# A standardized mosquito-to-mouse transmission assay with spatially defined bite areas

**DOI:** 10.64898/2026.11.19.767790

**Authors:** Elliott F. Miot, Hacène Medkour, Lauryne Pruvost, Louise Belleville, Mihra Tavadia, Félix Rey-Cadilhac, Carole Ginibre, Besthsabée Scheid, Pascal Boutinaud, Julien Pompon

**Affiliations:** MIVEGEC, Univ. Montpellier, IRD, CNRS, Montpellier, France

## Abstract

Mosquito-borne viruses initiate infection in the skin, where interactions between viral particles, host cells, and mosquito saliva can influence early infection and subsequent viral dissemination. Investigating these events requires methods that link mosquito exposure to the precise site of inoculation. However, identifying individual bite sites during subsequent tissue collection remains a practical challenge in mosquito-to-mouse transmission models. Here, we describe a standardized mosquito-to-mouse transmission assay that facilitates the localization of mosquito bite sites and targeted tissue sampling. The method combines a custom 3D-printed feeding device, individual mosquito exposure, and perforated adhesive tape to define the biting area and mark identified bite sites for subsequent collection. We demonstrate the application of this protocol using West Nile virus (WNV) and Dengue virus (DENV)-infected *Aedes aegypti* mosquitoes with C57BL/6J mice and A129 Ifnar1^-/-^ mice, respectively. All infected mosquitoes successfully blood-fed within 20 min of exposure. After three infected-mosquito bites, mice developed clinical signs from approximately day 6 post-bite, with mortality occurring between days 8 and 10 for WNV, while disease onset occurred at 4 days post bite and most of the mice recovered by day 10 for DENV. To investigate early WNV viral kinetics in skin, WNV genomic RNA was quantified in biopsies collected at marked bite sites and the draining inguinal lymph nodes. Viral RNA was detected in both tissues as early as 1 h post-bite, and RNA levels increased significantly at 24 h post-bite. This protocol provides a practical approach for identifying mosquito bite sites and investigating early tissue-level events following vector-mediated transmission. With appropriate validation, the protocol can be adapted to other vector–pathogen–host systems and downstream analyses.

**Summary:** Bite-initiated skin infection is a critical transmission bottleneck for arboviruses. A deeper understanding of these early events is essential for the development of intervention strategies. We describe a standardized mosquito-to-mouse transmission assay using a custom 3D-printed device and perforated tape to localize bite sites and investigate the early transmission steps.

## Introduction

Mosquito-borne viruses, including orthoflaviviruses [e.g., dengue virus (DENV), Zika virus (ZIKV) and West Nile virus (WNV)], alphaviruses (e.g., chikungunya virus) and orthobunyaviruses (e.g., Oropouche virus), represent an increasing global public health concern^1–3^. For all these pathogens, bite-initiated skin infection constitutes a critical step in the viral transmission cycle^4–7^. During probing, infected mosquitoes inoculate a small number of infectious virions together with saliva into the skin. The viruses subsequently establish infection in cutaneous structural cells, such as fibroblasts, and/or resident or recruited myeloid cells before disseminating to lymph nodes and peripheral organs, ultimately leading to systemic infection. The importance of these early events in the skin is supported by studies showing that modulation of cutaneous immune responses^8, 9^ or interference with salivary transmission-enhancing factors^6, 10–12^ can alter viral transmission efficiency and disease outcome. A deeper understanding of bite-initiated skin infection is essential for the development of intervention strategies targeting this critical transmission bottleneck^13, 14^.

Transmission assays require an *in vivo* model and have been established for several mosquito-borne viruses, although the susceptibility and suitability of the mouse model depend on the virus^15–17^. For WNV, immunocompetent adult mice, including C57BL/6J mice, develop neuroinvasive disease, with viremia, spread to the central nervous system and lethality that reproduce key features of severe human infection^18^. By contrast, DENV and ZIKV replicate poorly in immunocompetent mice, and productive infection requires interferon-deficient mouse model^19^. Compared with needle-based inoculation, mosquito-bite models preserve the contribution of mosquito saliva, probing behavior and the anatomical site of inoculation, providing a more physiologically-relevant model of arbovirus transmission. Current mosquito-to-mouse transmission models present several limitations: (i) variability in mosquito feeding efficiency; (ii) heterogeneity in the viral dose delivered during single bites or to each individual mouse; (iii) technical challenges associated with maintaining stable mosquito–mouse contact while minimizing animal disturbance; and (iv) limited precision in identifying and monitoring the exact bite site. Collectively, these constraints reduce experimental reproducibility and limit the utility of these models for investigating early cutaneous responses to mosquito-transmitted infection. Here, we describe a standardized mosquito-to-mouse transmission platform that enables precise localization of mosquito bite sites. Using this approach, we established a transmission assay with WNV and DENV, and characterized WNV early viral replication kinetics at the bite site.

## Protocol

### 1. Material preparation

1.1 Biting cage assembly

a. Using a 3D-printer, print the three components of the biting device from white PLA: the cover to maintaining the set of minicages, the set of 5 minicages to contain individual mosquitoes, and the base to hold the mouse. The device STL file is available in supplemental materials (**Figure 1A; Supplementary file 1)**.
b. Insert a laser-cut transparent PETG sheet through the slit in front of the set of minicages. Use a PETG sheet of 0.8 mm and dimensions of 55 x 26 mm. Secure the sheet in place using cyanoacrylate glue.
c. Attach mosquito mesh to the bottom of the set of minicages and secure it using cyanoacrylate glue.
d. Allow the glued components to dry completely before use.

1.2 Anesthesia

a. Prepare the ketamine/xylazine anesthesia cocktail at the concentrations specified in the approved animal-use protocol.
b. Dilute the cocktail with sterile saline or sterile water.
c. Prepare the required volume immediately before use and maintain it according to institutional guidelines.

1.3 Biting tape

a. Cut pieces of autoclave tape of 6 cm long
b. Using a 2 mm biopsy punch, perforate four holes per centimeters in a square pattern so they align with the minicage screened area using the 3D-printed template (**Supplementary file 2)**.
c. Tape them on a plastic sheet for later use.

NOTE: Because the mosquito proboscis penetrates < 2 mm into mouse skin in any direction^20^, we restricted the hole size to 2-mm-diameter.

**Figure 1.**
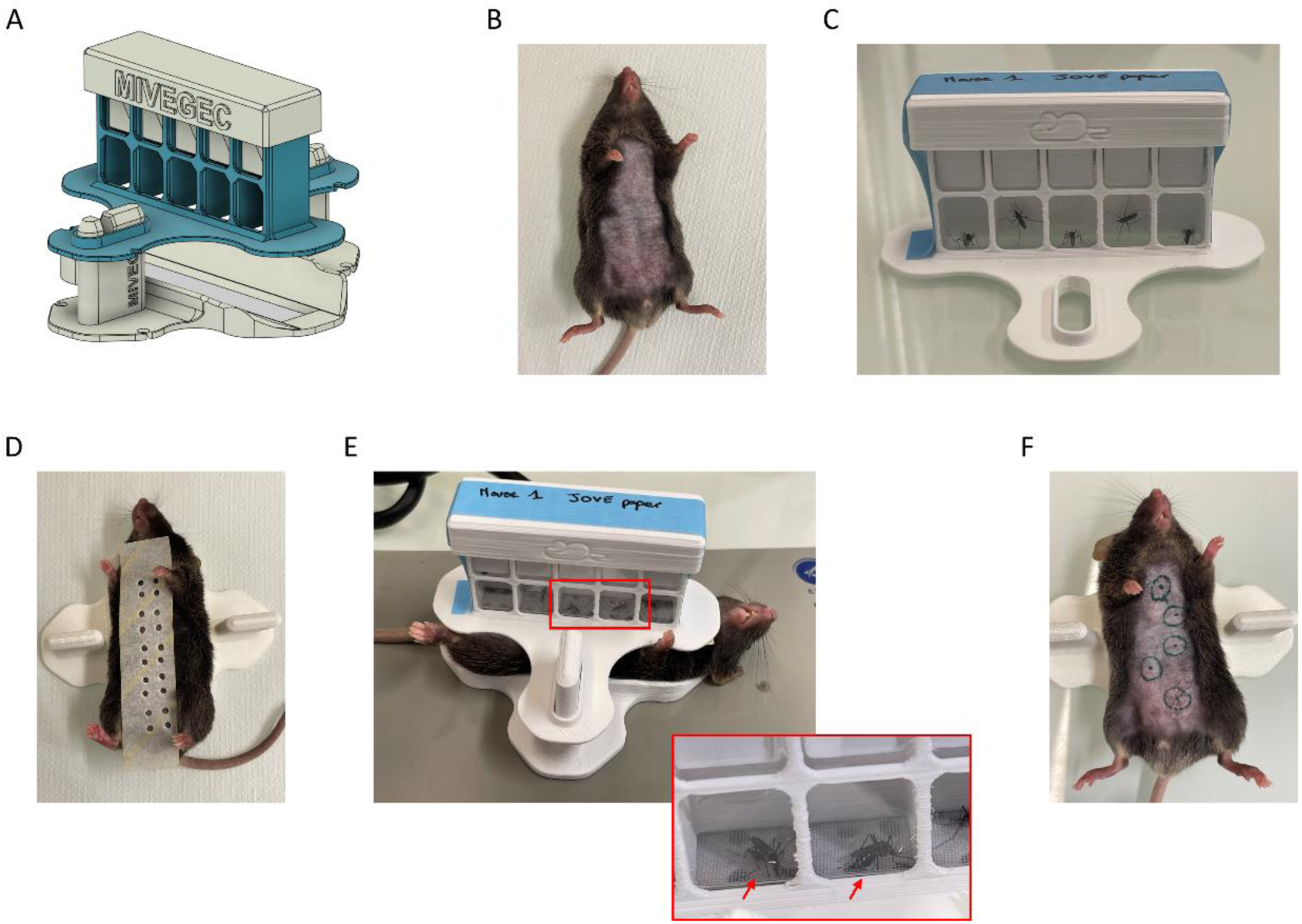
Experimental set-up. (**A**) 3D model of the biting device. (**B**) Shaved mouse ventral side. (**C**) A set of 5 minicages containing mosquitoes, secured with a taped cover. (**D**) Perforated tape positioned onto a mouse. (**E**) Complete experimental set-up during the biting phase. Arrows indicate bloodfed mosquitoes. (**F**) Marking of the bite sites on the mouse ventral side.

NOTE: Other punch size or number of holes can be used depending on your experimental setting. However, we observed that less holes reduces biting efficiency.

1.4 Mosquito collection tubes

a. Prepare and label 1.5 mL tubes with 1-mm diameter glass beads and TRK lysis buffer (EZNA total RNA kit I, Omega Bio-tek Inc.) to collect mosquitoes following the biting procedure.
b. Ensure that all tubes are appropriately identified.

1.5 Tissue collection tubes

a. Prepare and label 1.5 mL tubes with glass beads and TRK lysis buffer to collect animal tissues following the biopsy procedure. One tube per tissue sample.
b. Ensure that all tubes are appropriately identified.

### 2. Mosquito infection

1.6 Infect 3 to 7 day-old mosquitoes with the pathogen of interest using the appropriate validated infectious procedure (e.g., microinjection or infectious bloodmeal)^21, 22^.
1.7 Maintain infected mosquitoes in a controlled-environment chamber at 28°C and 80% relative humidity with a 10% sugar solution *ad libitum*.
1.8 Monitor mosquito survival during the incubation period.
1.9 At the appropriate time point following infection, proceed to the biting procedure described below.

## 3. Mosquito biting

NOTE: This protocol with infected mosquitoes must be realized in the containment level appropriate for the studied pathogen.

NOTE: Animal procedures have to be validated by the appropriate authority.

3.1 Preparation on the day before infection

a. Trim the ventral surface of each mouse with an electric trimmer (e.g., Battery-operated Trimmer Exacta, Aesculap Schermaschinen GmbH), extending from the region above genitalia towards the neck (**Figure 1B)**.

NOTE: Hairs can further be removed with shaving cream.

NOTE: For experiment assessing immune response, hairs should be removed one day before biting to avoid confounding effect of inflammation resulting from hair removal.

b. Remove the sugar solution from the mosquito cages 6 to 16 h before the planned biting procedure to promote host-seeking and blood-feeding behavior.

NOTE: Longer period of starvation reduces blood-feeding efficiency.

3.2 On the day of biting, transfer infected mosquitoes into minicages

a. Retrieve the mosquito cages from the controlled-environment chamber.
b. Cold-anesthetize the mosquitoes on ice for at least 10 min.
c. Inspect each cage and record the number of dead mosquitoes.
d. Separate live mosquitoes from dead ones.
e. Transfer one mosquito into each minicage.

NOTE: Always prepare additional mosquito-filled minicages in case some mosquitoes do not bite.

f. Secure the cover of the minicage set using tape on both sides (**Figure 1C)**.
g. Return the loaded minicages to the controlled-environment chamber and allow the mosquitoes to recover from cold anesthesia for at least 20 min.

NOTE: Avoid prolonged storage of mosquito-loaded minicages, as mosquito feeding efficiency will decrease over time. When a large number of mosquitoes are required, process the mosquitoes in batches rather than preparing all minicages simultaneously.

3.3 Mouse anesthesia

a. Restrain the mice using an appropriate handling technique. Ensure that the mouse is safely handled and immobilized.
b. Administer the anesthesia cocktail by intraperitoneal injection according to the approved animal-use protocol.
c. Return the mouse to its cage and allow sufficient time for anesthesia to take effect.
d. Confirm adequate anesthesia by gently pinching the rear foot for sign of a withdrawal response. If the mouse is still reacting, put it back in the cage and wait, otherwise, proceed with the experiment.
e. Continue to monitor the mouse throughout the procedure and maintain appropriate body temperature.

3.4 Mice exposure

3.4.1 Position the base of the biting device on the heating plates set at 37°C.
3.4.2 Place the anesthetized mice on each base with the dorsal surface in contact with the heating plate.
3.4.3 To define the biting area, place a piece of autoclave biting tape previously punctured on the shaved ventral skin (**Figure 1D**).
3.4.4 Position the minicage set over the mouse and align each minicage with the desired biting area (**Figure 1E**).
3.4.5 Monitor the mosquitoes during exposure and identify successful blood feeding by observing abdominal distension associated with blood intake.

NOTE: In the event that one or more mosquitoes do not bite in the first 5-10 min, switch the minicage with a new one.

3.4.6 Record the location of mosquito bloodfeeding in the experimental spreadsheet (**Supplementary file 3**).
3.4.7 Carefully remove the minicage set while keeping the autoclave tape in position.
3.4.8 Mark each identified biting site through the corresponding opening in the tape using a marker. Remove the tape. Circle the marks to facilitate subsequent identification of biopsy sites (**Figure 1F**).
3.4.9 Record the following information on the tape securing the cover of each biting cage:

3.4.9.1 Mouse identification number
3.4.9.2 Number of successfully engorged mosquitoes
3.4.9.3 Experiment identification number
3.4.10 Return each mouse to its respective cage.
3.4.11 Place the minicages in -70°C for at least 30 min to kill mosquitoes and transfer them to the prepared collection tubes for later processing.

## 4. Example of downstream application: Evaluation of skin and lymph node infections, and monitoring of disease outcome

4.1 Skin and draining lymph node biopsies

a. After the desired incubation period, anesthetize mice as described above.
b. Identify the previous marked mosquito bite sites.

NOTE: Marker-based identification remains visible for up to 24h. After this duration, the marker may fade. For longer incubation, use different method for skin bite identification.

c. Make a small incision below the bite site and pass a sterile grooved director under the skin so it is aligned under the marked site.
d. Position a sterile disposable 2 mm biopsy punch over the marked site.
e. Twist in both directions against the grooved director until it cuts through the skin and use sterile forceps and scissors to dissect the skin tissue circle.
f. Transfer the skin biopsy into a sterile 1.5 mL collection tube previously prepared.
g. Following completion of the skin collection, euthanize the mouse by cervical dislocation before recovery from anesthesia.
h. Dissect the inguinal draining lymph nodes (LN).
i. Transfer the LN into a sterile 1.5 mL tube previously prepared.
j. Process or store the collected tissues according to the downstream assay requirements.

4.2 Clinical monitoring

4.2.1. Monitor mice daily following infection.
4.2.2. Record body weight and clinical signs using the predefined scoring grid (**Supplementary file 4**).
4.2.3. Apply humane endpoints and intervention criteria to the animal-use protocol. The Supplementary file 4 includes automatically calculated humane endpoints.

## Representative Results

### 5. Mosquito-to-mouse WNV transmission model

Following this protocol, we established a controlled mosquito-to-mouse transmission model using WNV-infected *Ae. aegypti* mosquitoes with C57BL/6J mice and DENV-infected *Ae. aegypti* mosquitoes with A129 Ifnar1^-/-^mice.

Experiments were conducted in the VectoPole facility (MIVEGEC, Montpellier, France) which houses BSL1-3 insectary and animal facilities. *Ae. aegypti* BORA strain female mosquitoes^23^ were microinjected with 5 PFU (Plaque Forming Unit) of WNV strain IS98-ST1^24^ or 100 FFU (Focus Forming Unit) of DENV 2 strain D2Y98P^25^ using a nanoject (Drummond) and kept for 10 or 15 days post-injection, respectively. Each mouse was exposed to three bites from distinct mosquitoes following the above protocol. 100% of infected mosquitoes had blood fed within 20 min of exposure to mouse skin. A subset of the mosquitoes was tested by RT-qPCR to confirm proper infection. Mosquitoes contained an average of 1.5 x 10^9^ gRNA copies for WNV (**Figure 2A**) and 1.2 x 10^7^ copies for DENV (**Figure 2B**).

**Figure 2.**
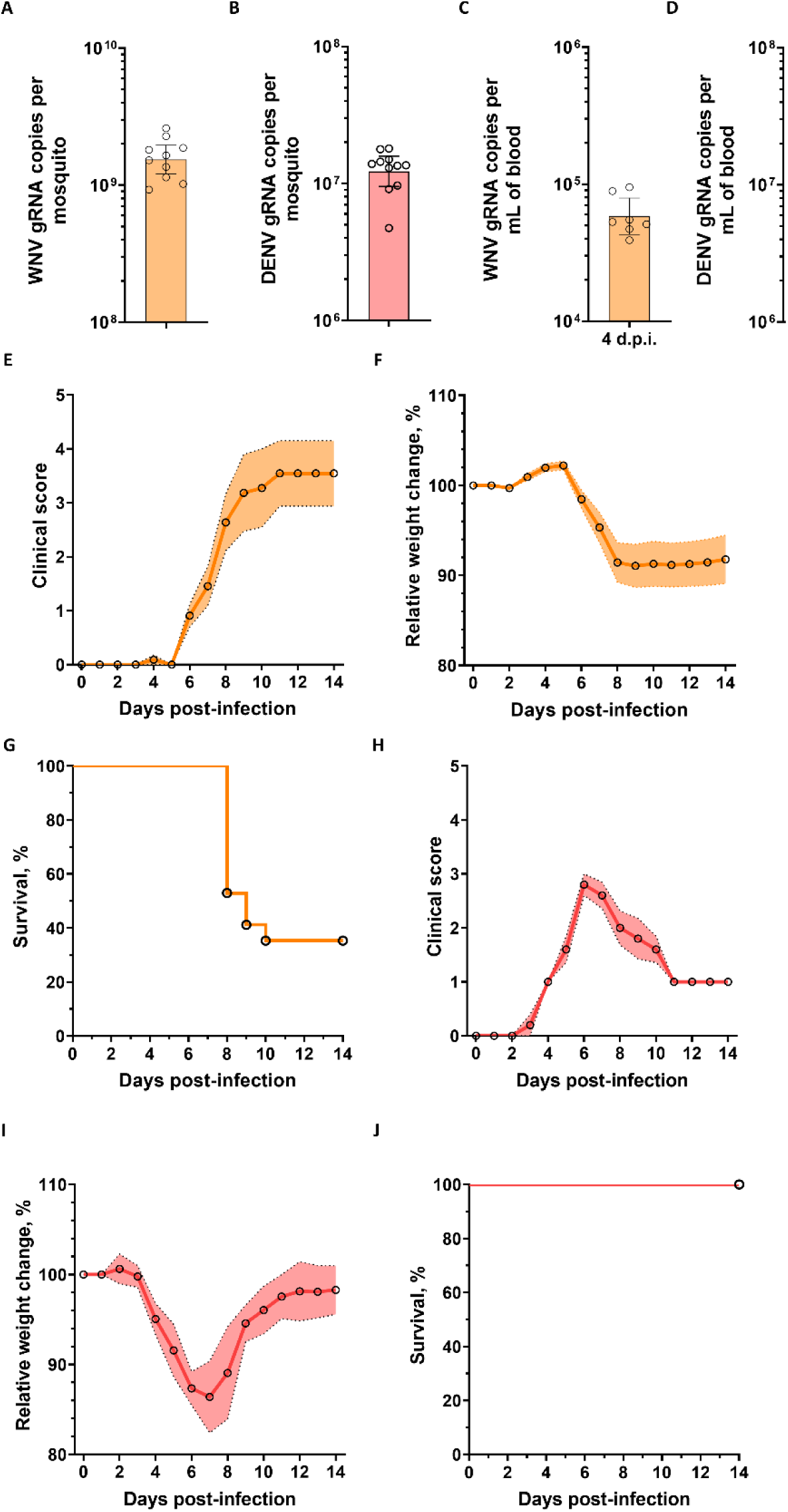
Disease outcome evaluation. (A,B) Geometric mean ± 95% CI gRNA copies per mosquito inoculated with WNV (A) and DENV (B). **(C,D)** Geometric mean ± 95% CI gRNA copies per mL of mouse blood collected at 4 days post bite with WNV-(C) and DENV-(D) infected mosquitoes. (**E-G**) Mean ± s.e.m. daily clinical signs (E), mean ± s.e.m. relative weight change (F), and survival rate following WNV-infected mosquito bite (G). N, 11 mice. (**H-J**) Mean ± s.e.m. daily clinical signs (H), mean ± s.e.m. relative weight change (I), and survival rate following DENV-infected mosquito bite (J). N, 5 mice. A-D Points indicate samples.

### 6. Assessment of disease outcome

Daily clinical sign evaluation using a score grid, body weight (**Table 1, Supplemental file 4**) and survival were recorded over 14 days post infectious bite. Successful transmission was verified by testing the gRNA level in mouse blood at the pic of RNAemia 4 days post infectious bite. RNAemia reached an average of 5.8 x 10^4^ gRNA copies per mL of blood for WNV (**Figure 2C**) and 7.5 x 10^6^ gRNA copies per mL of blood for DENV (**Figure 2D**). WNV disease onset including first clinical symptoms and bodyweight loss were observed six days post infectious bite (**Figure 2E, F**). Death occurred from day 8 until day 10 with a final survival rate of 35% (**Figure 2G**). Animals reaching predefined humane endpoints were euthanized and counted as events for survival analysis. DENV disease onset was observed four days post infectious bite and almost all mouse recovered by day 10 (**Figure 2H, I**). No mortality was observed (**Figure 2J**). These observations of symptoms confirm successful transmission.

**Table 1.** Example of a clinical grid score filled for a WNV-infected mouse. Clinical scoring [0-3] with 0 = normal, 1 = mild, 2 = moderate and 3 = severe.

| **Mouse 1** | **Day 0** | **Day 1** | **Day 2** | **Day 3** | **Day 4** | **Day 5** | **Day 6** | **Day 7** | **Day 8** | **Day 9** |
|---|---|---|---|---|---|---|---|---|---|---|
| Weight (g) | 28.6 | 28.6 | 28.1 | 28.3 | 28.9 | 29 | 27.6 | 26.4 | 24.7 | 23.2 |
| % Weightloss (g) vs Max weight | 0.00 | 0.00 | 1.75 | 1.05 | 0.00 | 0.00 | 4.83 | 8.97 | 14.83 | 20.00 |
| Weightloss score (see Rules in suppl. file 3) | 0 | 0 | 0 | 0 | 0 | 0 | 1 | 1 | 2 | 3 |
| Weakening of the hind limbs [0-3] |  |  |  |  |  |  |  | 1 | 1 | 2 |
| Hunched posture [0-3] |  |  |  |  |  |  | 1 | 2 | 2 | 2 |
| Rough or unkempt fur [0-3] |  |  |  |  |  |  |  | 1 | 2 | 2 |
| Decreased activity [0-3] |  |  |  |  |  |  |  |  | 1 | 1 |
| Labored breathing [0-3] |  |  |  |  |  |  |  |  |  |  |
| Decrease in body condition score (protruding bones, especially in the hindquarters) [0-3] |  |  |  |  |  |  |  |  |  |  |
| Neurological signs [0-3] |  |  |  |  |  |  |  |  |  |  |
| **Death overnight (yes=1)** |  |  |  |  |  |  |  |  |  |  |
| **Extended score** | 0 | 0 | 0 | 0 | 0 | 0 | 2 | 5 | 8 | 10 |
| **Reduced score** | 0 | 0 | 0 | 0 | 0 | 0 | 1 | 2 | 3 | 4 |
| **Mortality (see Rules in suppl. file 3)** |  |  |  |  |  |  |  |  |  | **x.wl** |
| Blood sampling (µL) |  |  |  |  | 20 |  |  |  |  |  |

### 7. Early WNV kinetics at the bite site and draining lymph nodes

At 1-, 6-, 9-, and 24-h post mosquito bites, we collected skin biopsies of the bitten areas and the pair of draining inguinal LN. WNV genomic RNA (gRNA) copies were quantified by RT-qPCR as in Medkour et al. 2025^6^. We first calculated the infection rate in skin as the percentage of positive RT-qPCR samples over the total number of samples. The infection rate ranged from 55% to 72% with no significant difference between time points (**Figure 3A)**. Second, we quantified the gRNA copies in positive samples (i.e., those in which at least one copy of gRNA was detected). At 1h post bite, corresponding to the delivered inoculum, skin samples contained an average of 1.55 x 10^2^ gRNA copies (**Figure 3B)**. Whereas the skin infection level remained constant until 9h post bite, it significantly increased to 1.99 x 10^3^ gRNA copies per skin sample at 24h post-bite. We similarly reported rate and level of infection in LN. As early as 1h and until 6h post bite, 100% of LN had detectable copies of gRNA. However, at 9h post bite, infection rate decreased to 83%, before resuming to 100% at 24h post bite (**Figure 3C**). Infection level was constant until 9h post bite and then significantly increased at 24h post bite (**Figure 3D**). Together with our previous studies showing a positive correlation between bite-initiated skin infection on LN infection^6, 10^, these results are coherent with a model in which LN infection results from a second wave of viral particles released during active viral replication in the skin.

**Figure 3.**
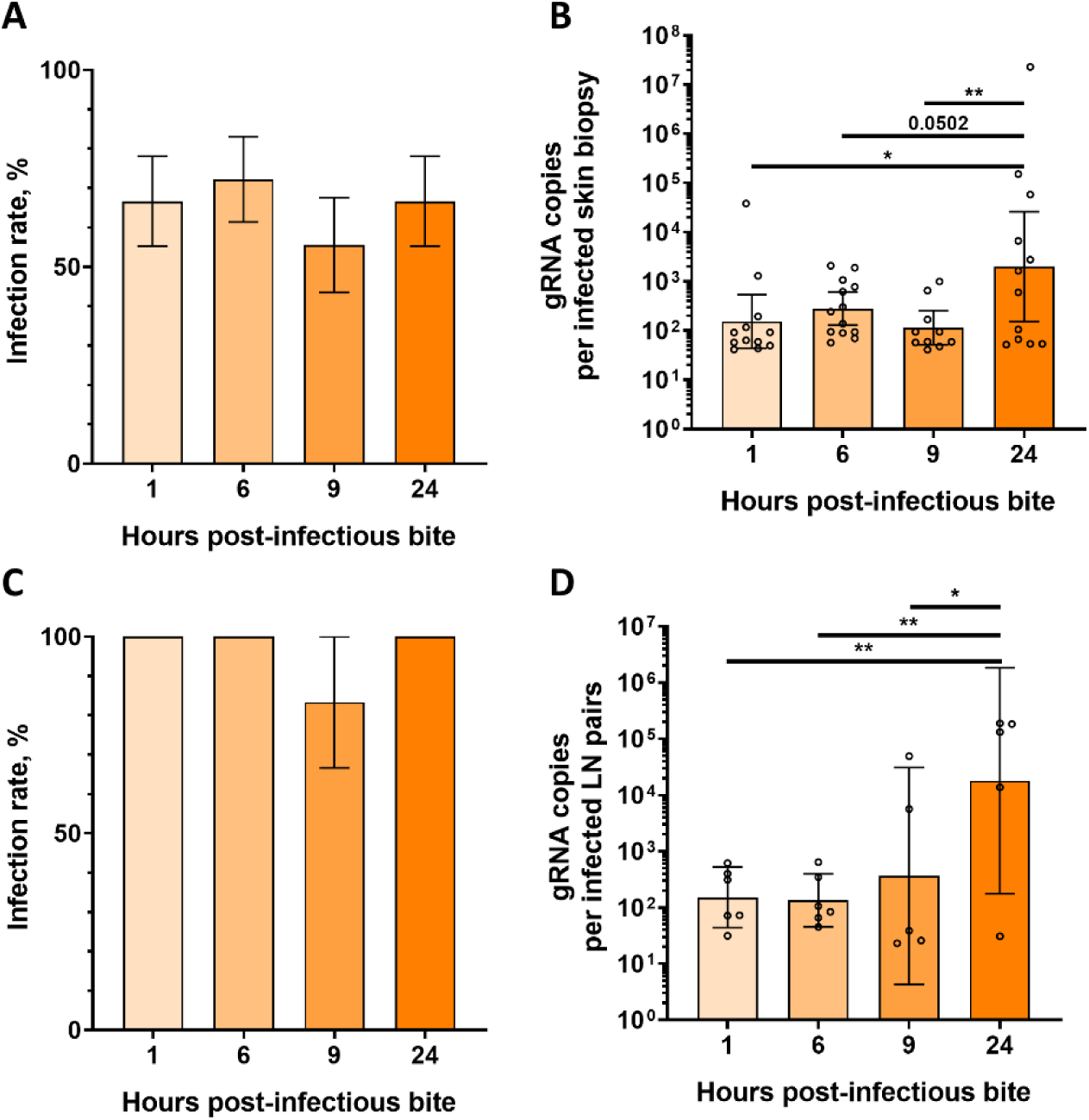
Early WNV infection kinetics at the bite site and in draining lymph nodes. WNV infection was quantified at 1, 6, 9, and 24h post-bite. **(A, B)** Infection rate (A) and infection level (B) in skin samples. **(C, D)** Infection rate (C) and infection level (D) in pairs of draining LNs. Infection rate was defined as the number of positive samples over the total number of analyzed samples. Infection level was calculated from the positive samples. A,C Bars represent proportion ± standard error. B, D Bars represent geometric mean ± 95% C.I. Points indicate samples. Multiple comparisons were performed using Fisher’s LSD post-hoc analysis. * *p*<0.05; ** *p*<0.01. N mice, 6. N skin sample per mouse, 3.

## Discussion

We developed a standardized mosquito-to-mouse transmission assay that enables precise localization of mosquito bite sites for the investigation of early arbovirus skin infection. The protocol combines optimized mosquito preparation, individual mosquito exposure, and a custom 3D-printed feeding device with perforated adhesive tape to define biting areas and facilitate subsequent tissue sampling. Using both WNV-infected *Ae. aegypti* mosquitoes and C57BL/6J mice and DENV-infected *Ae. aegypti* mosquitoes and Ifnar1^-/-^ mice as a proof of concept, we demonstrate the feasibility of this approach for studying disease outcome and the temporal distribution of viral RNA in the skin and draining LN. By allowing the identification and targeted sampling of the bite sites, this method addresses an important limitation of conventional mosquito-bite models, in which the precise location of viral inoculation can be difficult to determine. This approach provides a practical experimental platform for investigating the early events following mosquito-mediated transmission and the local host–virus interactions that contribute to arbovirus skin infection and dissemination. The overall biting device is easy to set-up, cost-effective and offers a standardized transmission assay compatible with diverse downstream applications.

Mosquito-mediated transmission initiates infection in skin, where viral particles interact with structural and immune cells in the presence of mosquito saliva, which can modulate local immune responses and influence viral dissemination^7^. Mosquito saliva and bite-induced inflammation have been reported to modulate local immune responses and influence arbovirus infection and pathogenesis^26, 27^. These findings highlight the importance of investigating the local environment at the bite site. However, experimental investigation of the early events of skin infection requires precise localization of the mosquito bite. By defining the biting area and facilitating its subsequent identification, our method provides a practical means for linking mosquito exposure to targeted tissue sampling. This spatial approach may be particularly useful for investigating local immune responses, viral dissemination, and interactions between mosquito saliva and host tissues during the early stages of transmission.

Using the method, we detected WNV gRNA in both skin and draining lymph nodes as early as 1 h post-bite. Interestingly, while WNV gRNA levels remained constant in the skin and LN during the first 24 h, it increased significantly in both tissues at 24 h post-bite. This observation is consistent with the model of progressive viral dissemination from skin to draining LN, as shown in our previous work^9^.

Beyond its application to WNV and DENV transmission, this protocol provides a flexible platform for investigating early host–virus interactions at the bite site for a diverse range of vector-pathogen system. The procedure can be adapted to different hematophagous insect vectors, such other mosquito species, Culicoides, and a variety of pathogens in a multitude of hosts. Variation in sampling time points and downstream analytical approaches can further help address a large range of questions. The model should, however, be empirically validated for each pathogen–vector-host system. Overall, this method provides a practical foundation for investigating the early stages of vector-borne transmission.

Several limitations to our approach should be considered. Although the protocol standardizes mosquito preparation and feeding conditions, variability in the quantity of infectious virus delivered during individual bites remains an inherent feature of vector-mediated inoculum delivery. This intrinsic biological variability in the amount of saliva released in the skin can be partially addressed by increasing the number of bites to normalize the dose delivered. Multiple bites do not alter local cutaneous viral load at 24 h post-bite but increase the viral inoculum reaching draining LNs^9^. The visibility of the bite-site marking may fade beyond 24 h, limiting the identification of sites in experiments requiring later sampling. For longer experiments, the persistence of the marking and the feasibility of re-marking should be evaluated for the specific experimental conditions.

## Acknowledgements

We thank T. Pellissier and P. de Féligonde from the company PSL3D (Montpellier, France) for their help and advice to create the 3D-models. We thank P. Desprès for providing the WNV strain and S. Alonso for the DENV strain.

## Disclosures

The authors declare that they have no conflicts of interest.

**Supplementary file 1. STL file of the biting device.**

**Supplementary file 2. STL file of the biting tape template.**

**Supplementary file 3. Spreadsheet to record bite sites.**

**Supplementary file 4. Clinical scoring grid.**

