## Supplementary figures and images for "A standardized mosquito-to-mouse transmission assay with spatially defined bite areas"

### Supplementary file 3

#### Mouse 1

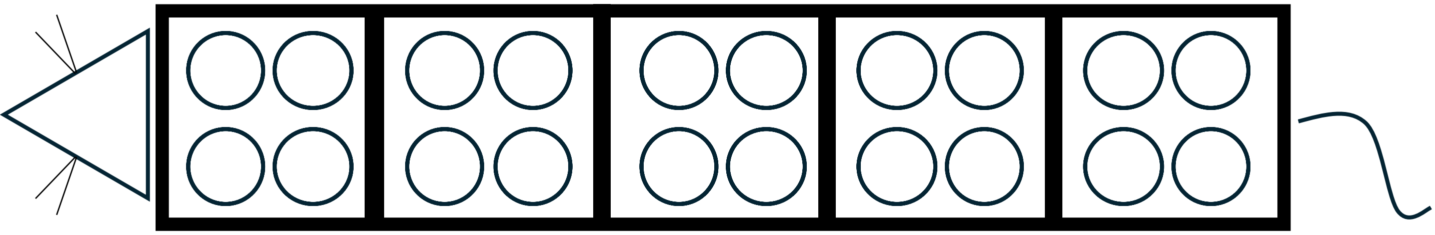

Bite time ............ ............ ............ ............ ............

#### Mouse 2

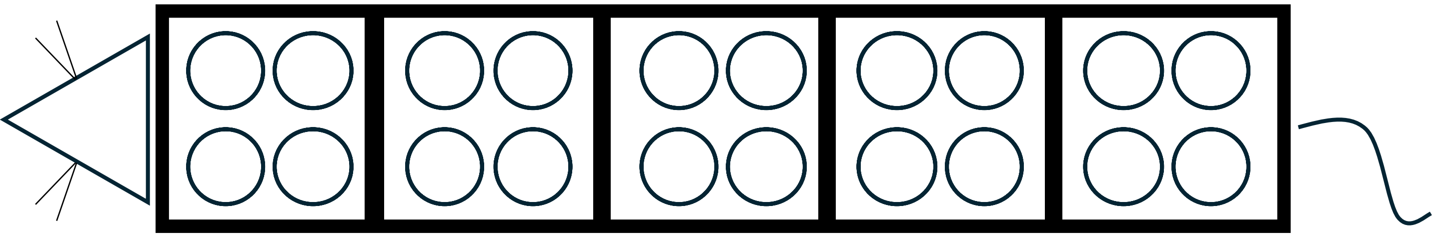

Bite time ............ ............ ............ ............ ............

#### Mouse 3

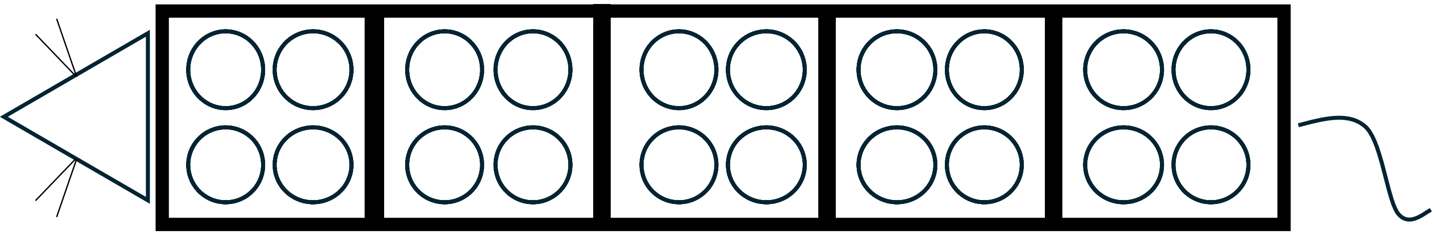

Bite time ............ ............ ............ ............ ............

#### Mouse 4

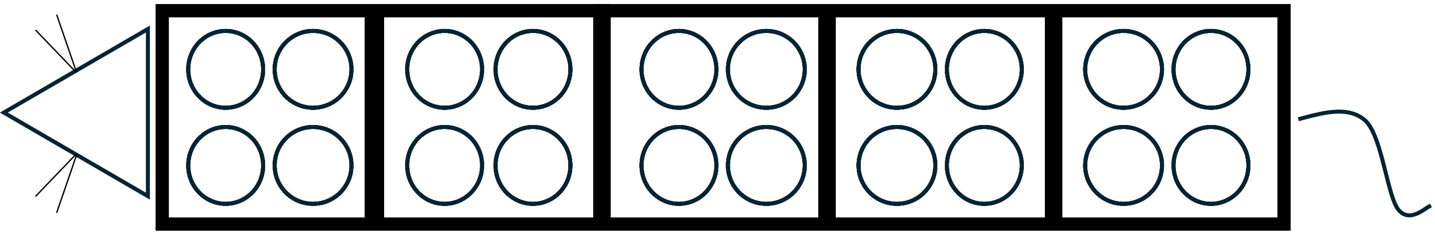

Bite time ............ ............ ............ ............ ............

#### Mouse 5

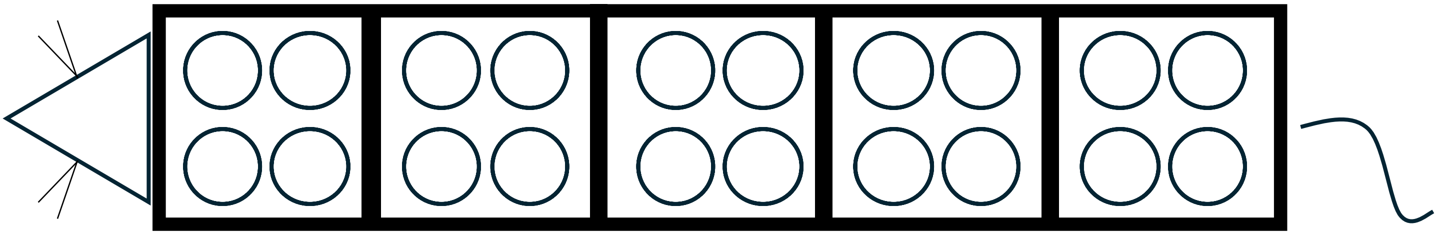

Bite time ............ ............ ............ ............ ............
